# Global chromatin mobility induced by a DSB is dictated by chromosomal conformation and defines the outcome of Homologous Recombination

**DOI:** 10.1101/2022.02.03.478935

**Authors:** Fabiola Garcia Fernandez, Etienne Almayrac, Ànnia Carré Simon, Renaud Batrin, Yasmine Khalil, Michel Boissac, Emmanuelle Fabre

## Abstract

Repair of DNA double-strand breaks (DSBs) is crucial for genome integrity. A conserved response to DSBs is an increase in chromatin mobility that can be local, at the site of the DSB, or global, at undamaged regions of the genome. Here we address the function of global chromatin mobility during homologous recombination (HR) of a single, targeted, controlled DSB. We set up a system that tracks HR *in vivo* over time and show that two types of DSB-induced global chromatin mobility are involved in HR, depending on the position of the DSB. In spatial proximal regions such as the pericentromeric region, a DSB therein induces global mobility that depends solely on H2A(X) phosphorylation and accelerates repair kinetics, but is not essential. In contrast, the global mobility induced by a DSB away from the centromere, becomes essential for HR repair and is triggered by homology search through a mechanism that depends on H2A(X) phosphorylation, checkpoint progression and Rad51. Our data demonstrate that global mobility is governed by chromosomal conformation and differentially coordinates repair by HR.

## Introduction

Double-strand breaks (DSBs) are the most harmful lesions that constantly challenge the eukaryotic genome, potentially leading to deleterious genetic alterations that include local modifications of the damaged DNA (insertions/deletions) or global chromosomal rearrangements (translocations). To faithfully maintain genome stability and survive DSBs, cells display a DNA damage response (DDR) involving complex networks that detect, signal and repair DSBs. Two of the most conserved factors contributing to DSB signaling are the checkpoint kinases Tel1 and Mec1 (mammalian ATM/ATR orthologues) that catalyze histone H2A phosphorylation in yeast and the mammalian variant H2AX, termed ƴ-H2A(X) (Waterman et al. 2020; Jackson and Bartek 2009). ƴ-H2A(X) extends widely on both sides of the DSB, up to 100 kb in yeast and 1 MB in mammals (Caron et al. 2012; Aymard et al. 2017; Rogakou et al. 1999; Iacovoni et al. 2010; Shroff et al. 2004). This wide spreading of ƴ-H2A(X) serves as a docking site for further DDR proteins such as the mediator protein Rad9 (53BP1 in mammals) (Kinner et al. 2008).

DSBs can be accurately repaired by homologous recombination (HR). This mode of repair, predominant in S phase, uses the sister chromatid as template (Symington and Gautier 2011). When homologous sequences are not found in the close vicinity of the break, a global genomic homology search may occur. Homology search starts with the formation of a presynaptic nucleoprotein filament that is composed of 3’-ssDNA overhangs coated with the recombinase protein Rad51 (Sugawara et al. 2003; Wang and Haber 2004; Chen et al. 2008; Kalocsay et al. 2009). The Rad51-nucleofilament enables the sampling and recognition of homology within the nucleus through base pairing (reviewed in (Candelli et al. 2014). By analyzing Rad51 distribution in haploid yeast after the induction of a single DSB by Chromatin Immunoprecipitation (ChIP), a study visualized ‘snapshots’ of on-going homology search, and found Rad51 strikingly distributed over a large portion of the broken chromosome (Renkawitz et al. 2013b). Notably, the genome-wide Rad51 signal distribution was concomitant with that of Mec1- dependent ƴ-H2A(X), leading to the conclusion that Mec1 might be attached on the probing end during homology search. The resulting phosphorylation of H2A(X) could thus promote chromatin remodeling, signaling or repair.

The non-random organization of chromosomes in the nucleus affects numerous nuclear processes including homology search. In the configuration of budding yeast chromosomes (known as the Rabl configuration), centromeres are clustered close to the yeast microtubule organizing center (Spindle Pole Body), as evidenced by abundant *trans*-contacts between centromeres, and telomeres are bound to the nuclear periphery in a position that depends on the size of the chromosome arms (Jin et al. 2000; Duan et al. 2010; Schober et al. 2008; Therizols et al. 2010). It was shown in yeast that DSBs recombine more efficiently in spatially proximal regions (Agmon et al. 2013; Batté et al. 2017; Lee et al. 2015). For example, recombination between homologous sequences that lie close to centromeres is more efficient than between regions that are distantly located on chromosome arms (Agmon et al. 2013). Accordingly, induction of a DSB close to a centromere triggers concomitant Rad51 and ƴ-H2A(X) signals within centromeric regions of all other yeast chromosomes, suggesting that homology search can efficiently occur on DNA that is located proximal to the centromere of other chromosomes (Renkawitz et al. 2013b; Lee et al. 2014). Likewise, mammalian topologically associated domains (TADs), where chromatin contacts are enriched, promote diffusion of ATM, which allows DSB signaling and thus repair (Arnould et al. 2021).

The search for distantly located homologies challenges recombination, likely requiring additional sophisticated mechanisms, such as chromatin mobility. A conserved response to DSBs, initially observed in yeast, consists of increased chromatin mobility both at the damaged locus and elsewhere in the undamaged genome, referred to as local and global mobility, respectively (reviewed in (Fabiola García Fernández 2022; Seeber et al. 2018; Oshidari et al. 2020; Smith and Rothstein 2017). Different mechanisms have been described to explain these DSB-induced chromatin dynamics, ranging from intrinsic changes in chromatin properties, including chromatin stiffening or decompaction, to extrinsic nuclear organizing factors such as actin and microtubules (Herbert et al. 2017; Miné-Hattab et al. 2017; Hauer et al. 2017; Lawrimore et al. 2017; Strecker et al. 2016; Caridi et al. 2018). Functionally, the mobility of damaged and undamaged chromatin has been implicated in various DDR processes such as DSB relocation, clustering, and homology search. Indeed, several studies have proposed that both mobilities allow homology probing in a larger nuclear volume (Mine-Hattab and Rothstein 2012; Dion et al. 2012; Seeber et al. 2013; Smith et al. 2018). Accordingly, given the concomitant spreading of Rad51 and ƴ-H2A(X) proposed to take place during homology search, the increase in local and global mobility is abolished in the absence of Rad51 or ƴ-H2A(X) (Mine-Hattab and Rothstein 2012; Miné-Hattab et al. 2017; Smith et al. 2018; Garcia Fernandez et al. 2021). However, the role of global mobility in HR remains to be clarified. Indeed, both a positive correlation between global mobility and recombinant products formation, as well as a lack of effect of global mobility on recombination are documented (Challa et al. 2021; Cheblal et al. 2020; Hauer et al. 2017; Mine-Hattab and Rothstein 2012; Miné-Hattab et al. 2017; Dion et al. 2012; Strecker et al. 2016). Similarly, mobility within a mammalian TAD, allowing ATM to reach other regions of chromatin, has also been suggested to explain the favorable environment for HR observed within a TAD (Schrank et al. 2018; Aymard et al. 2017). Although the mechanisms and functions of increased mobility are not fully understood, it is now considered part of the DDR.

To characterize the role of global chromatin mobility upon DNA damage during HR in yeast, we set up a system in haploid cells that allows simultaneously tracking HR at the single cell level and following global chromatin dynamics upon a single DSB induction, taking into account chromosome conformation. We show that the global increase in mobility induced by a DSB is influenced by the conformation of the chromosome. In the pericentromeric region, a DSB close to the centromere induces a global mobility that we call ‘proximal’. This mobility that occurs *in trans* towards the DSB is dependent on ƴ-H2A(X), but neither on the Rad9-dependent checkpoint nor on the Rad51 nucleofilament. Therefore, this ‘proximal’ mobility has little influence on the overall HR efficiency, but rather accelerates repair kinetics. In contrast, a subtelomeric DSB induces a mobility that we call ‘distal’. This distal mobility only occurs if a homologous sequence is present and if it is near the centromere. We demonstrate that the donor homologous sequence allows ƴ-H2A(X) spreading that is the requisite for chromatin motion. This distal mobility shares the characteristics of homology search mobility. It is indeed dependent on Rad9 and the Rad51 nucleofilament, and is necessary for homologous recombination. Our data unambiguously demonstrate the role of damage-induced global mobility due to non-random chromosome organization and unify conflicting data on its role in homologous recombination repair.

## Results

### THRIV, a system to Track Homologous Recombination In Vivo

We have recently shown that chromatin mobility favors NHEJ repair in the absence of a donor sequence (Garcia Fernandez et al. 2021). Here, we asked whether global mobility has a role on DSB repair when a homologous donor sequence is present. To answer this question, we designed a system that simultaneously tracks HR at the single-cell level and measures global genome mobility away from the DSB. We chose to follow HR through fluorescent protein *mCherry* synthesis, which coding sequence was used as both recipient and donor sequences. Neither of these sequences was able to translate into a functional fluorescent protein, unless HR occurred. The *mCherry* recipient sequence was chromosomal under the control of the TEF constitutive promoter and contained an I-*Sce*I cutting site (cs), followed by a stop codon that truncates mCherry. The donor sequence, carried by a centromeric plasmid, hereafter referred to as dCen, consisted into an *mCherry* CDS lacking promoter and terminator sequences, sharing 539pb and 172pb of homology upstream and downstream the I-*Sce*I cs of the recipient sequence, respectively (**Fig. 1A**). The *mCherry* recipient sequence was integrated at different positions: at 5kb from the centromere of chromosome IV, generating the so-called C strain; in a luminal position at respectively 400kb and 1060kb from the centromere and the telomere (L strain) and a subtelomeric position at 10kb from the right telomere (S strain) (**Fig. 1B**). All strains grew with a similar division time **(Supp. Fig 1A).** These different recipient sequence positions, near and far from the characteristic anchoring features of yeast chromosomes, allowed us to consider distinct chromosome architecture contexts.

**Figure 1.**
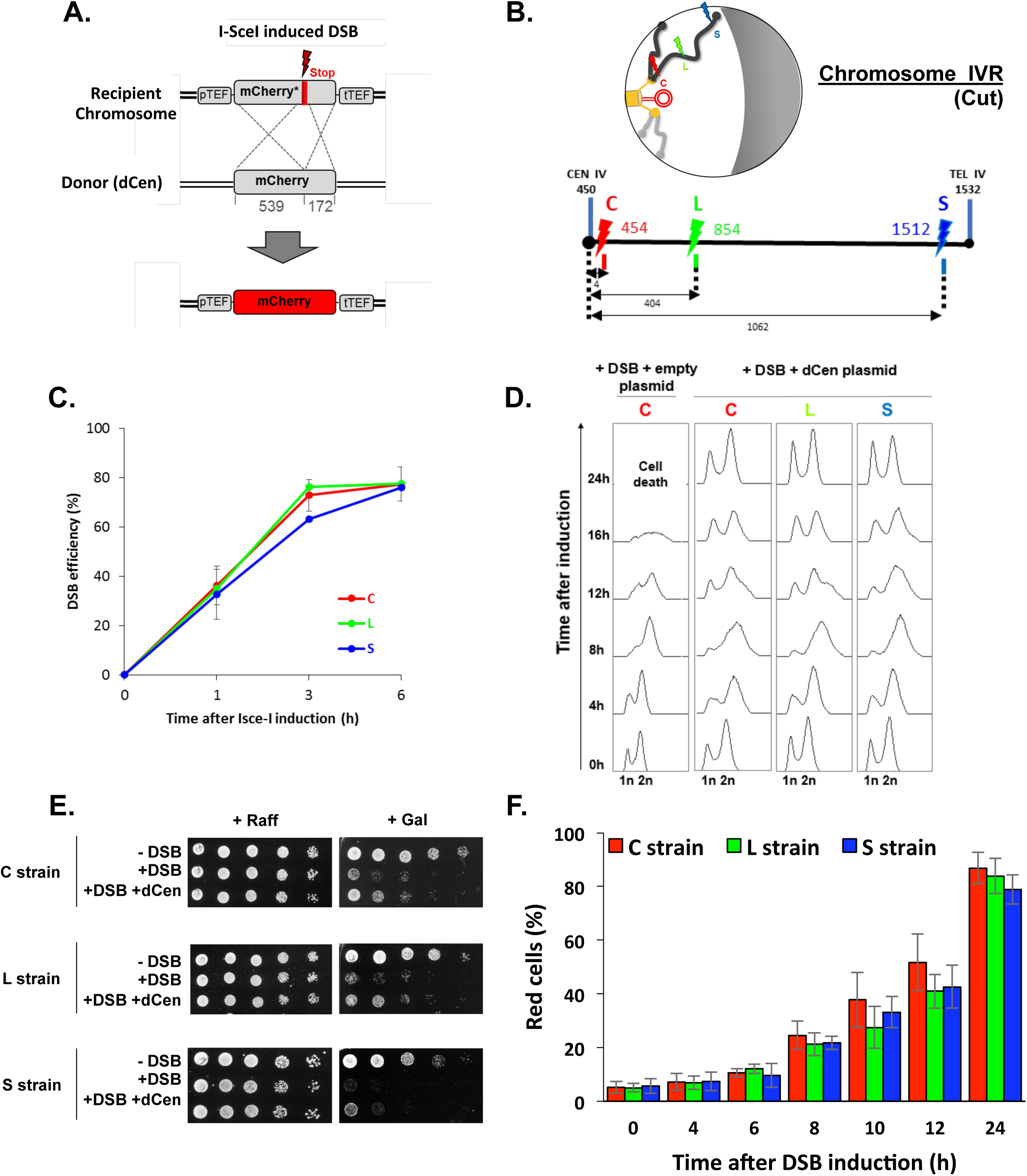
THRIV, a system to follow homologous recombination (HR) in vivo. **A**. Schematic representation of the two sequences used to track recombination events during time. The recipient *mCherry-I-SceI* has a 30pb I-*Sce*I sequence followed by a stop codon (red bar) inserted at 57 amino acids from the end of the *mCherry* coding sequence. The donor (dCen) is a full-length *mCherry* lacking promoter and terminator sequences, carried on a centromeric plasmid. A functional mCherry could be expressed only after HR. **B**. Schematic 2-D representation of the Rabl spatial configuration of two chromosomes in the nucleus of *Saccharomyces cerevisiae*. The Spindle Pole Body (rectangle), the microtubules (lines) and the centromeres (spheres) are in yellow. Telomeres are shown as grey/black spheres attached to the nuclear envelope. An orange circle represents the dCen plasmid. The grey crescent symbolizes the nucleolus. Lightening symbolize the targeted I-*Sce*I cutting sites (cs) on Chromosome IVR arm, with same color code as in the linear representation represented below. The cs are respectively located at 5kb from centromere CEN IV (red, strain C), 400kb from CEN IV (green, strain L) and 20kb from telomere TEL IVR (blue, strain S), distances are in kb. **C**. I-*Sce*I cleavage efficiency is measured in a non-donor strain by qPCR using primers flanking the I-*Sce*I cs. The error bars represent the standard deviation of three independent experiments. **D**. FACS analysis of the cell cycle of strains C, L and S after induction of I-*Sce*I with or without dCen plasmid. For the quantification of the cell cycle progression, the amount of DNA is measured with Sytox after fixation with ethanol and treatment with RNase. The peaks corresponding to 1n and 2n amount of DNA are indicated. **E**. Drop assay (tenfold serial dilutions) showing the sensitivity of C, L and S strains, containing a plasmid expressing or not I-*Sce*I (-DSB, +DSB respectively), with dCen plasmid or without (empty plasmid), when I-*Sce*I is induced (Galactose) or not (Raffinose). **F**. Homologous Recombination (HR) kinetics upon induction of I-*Sce*I in the presence of dCen in C, L or S strains. Percentages of repaired red cells were measured by FACS after PFA fixation in the absence of Sytox. Error bars represent the standard deviation of three independent experiments.

Expression of the I-*Sce*I endonuclease on a plasmid under the control of the *GAL1* galactose-inducible promoter generated single DSB (+DSB), whereas the empty plasmid not expressing I-*Sce*I was used as a control (-DSB). I*-Sce*I expression, measured by RT-qPCR showed a 6.7 ± 0.85 fold increase after 1h and a 5-fold plateau at longer times (**Supp. Fig. 1B**). Cutting efficiency, measured by qPCRs with primers surrounding the I-*Sce*I cs, showed 36%±7.8 at 1h and reached 77%±0.8 after 6h I-*Sce*I induction (**Fig. 1C**), indicating a delay between the enzyme expression and the effective cutting. This delay is not observed in the W303 background (Batté et al. 2017) and could be due to the BY background used here. Longer induction times of I-*Sce*I triggered cell cycle arrest and a low number of colonies visible by serial dilutions, independently of all chromosomal positions of the DSB (**Fig. 1D, 1E**). The presence of dCen plasmid resulted in both, the resolution of 12 hours cell cycle arrest and recovery of survival after DSB induction, as shown by FACS analyses and serial dilutions (**Fig. 1D, 1E**). Expression of mCherry was confirmed by fluorescent microscopy, suggesting that repair occurred by HR (**Supp. Fig1C**). The absence of I-*Sce*I cleavage *in vitro* in DNA extracted from red cells further established HR (**Supp. Fig. 1C**). Red cells were detected by FACS 4h after I-*Sce*I induction and their number, which included repaired cells and their progeny, progressively increased with time to 87.08 ± 4.43, 77.40 ± 0.54 and 72.58 ± 0.42% at 24h in C, L and S strains, respectively (**Fig. 1F**). The lower number of red cells in S strain, related to the slightly lower survival shown in serial dilutions, could be explained by the distant spatial position between the DSB near the telomere and the donor sequence, since the nuclear localization of the dCen plasmid indicated a position comparable to that of a pericentromeric sequence (**Supp. Fig. 1D**). Thus, after a single DSB, the system that <u>t</u>racks <u>h</u>omologous <u>r</u>ecombination in <u>v</u>ivo (THRIV) is efficient and depends on the relative spatial position between the donor and recipient sequences.

### Global chromatin mobility is enhanced during homologous recombination

A positive correlation between global mobility and the ability to repair a DSB by HR has been established (Challa et al. 2021; Cheblal et al. 2020; Hauer et al. 2017; Mine-Hattab and Rothstein 2012; Miné-Hattab et al. 2017; Dion et al. 2012), but it has also been shown that global mobility does not necessarily have a role in HR efficiency (Strecker et al. 2016). To directly investigate global mobility during HR, we tracked TetO repeats inserted at 96kb from CenV and bound by the tetR-GFP fusion protein (named V-Vis, for <u>vis</u>ualization of the locus on chromosome V, (**Fig. 2A** and (Strecker et al. 2016)), in strains harboring the THRIV system, i.e I-*Sce*I cutting sites at different positions and the donor sequence on a plasmid. We analyzed the mean squared displacements (MSDs) of V-Vis upon induction by I-*Sce*I in the different C, L and S strains, in the presence of the plasmids carrying or not the donor sequence. For each condition, we tracked the locus by time-lapse microscopy at 100ms time intervals in 500-1000 cells, as in (Herbert et al. 2017), and observed mobility for short (10s) and long (180s) time scales (**Fig. 2, Table S1 and Supp. Fig 2A**). Strains expressing or not I-*Sce*I were grown in galactose for 6h. In the absence of I-*Sce*I induction, the V-Vis locus exhibited a subdiffusive behavior, with MSDs exhibiting a power law (MSD ∼ Dtα), where α, was ∼0,6. The value of this anomalous exponent below 1 was previously shown at these short and long time scales for many loci scattered along the genome (**Fig. 2B, -DSB, Table S1** and **Supp. Fig. 2A**; (Spichal et al. 2016; Strecker et al. 2016; Herbert et al. 2017; Hajjoul et al. 2013; Dion et al. 2012; Mine-Hattab and Rothstein 2012; Seeber et al. 2013). Upon I-*Sce*I induction and generation of a single DSB in the C strain, the V-Vis locus showed an increased dynamic at the short time scale as well as an increased anomalous α, indicating changes in chromatin properties around the tracked locus (**Fig. 2B, +DSB, Table S1** and **Supp. Fig. 2B**)). Global dynamics after 6h-induction of I-*Sce*I was also observed in strain C at the longer time scale of 180ms (**Supp. Fig. 2A**) and gradually increased to a 1.95 fold after 12h (**Supp. Fig. 2B)**. In contrast, a single DSB (verified by qPCR, **Fig 1C**) in L and S strains had no effect on V- Vis dynamics, which remained similar to that observed in the absence of DSB induction (**Fig. 2B** **+DSB**).

**Figure 2.**
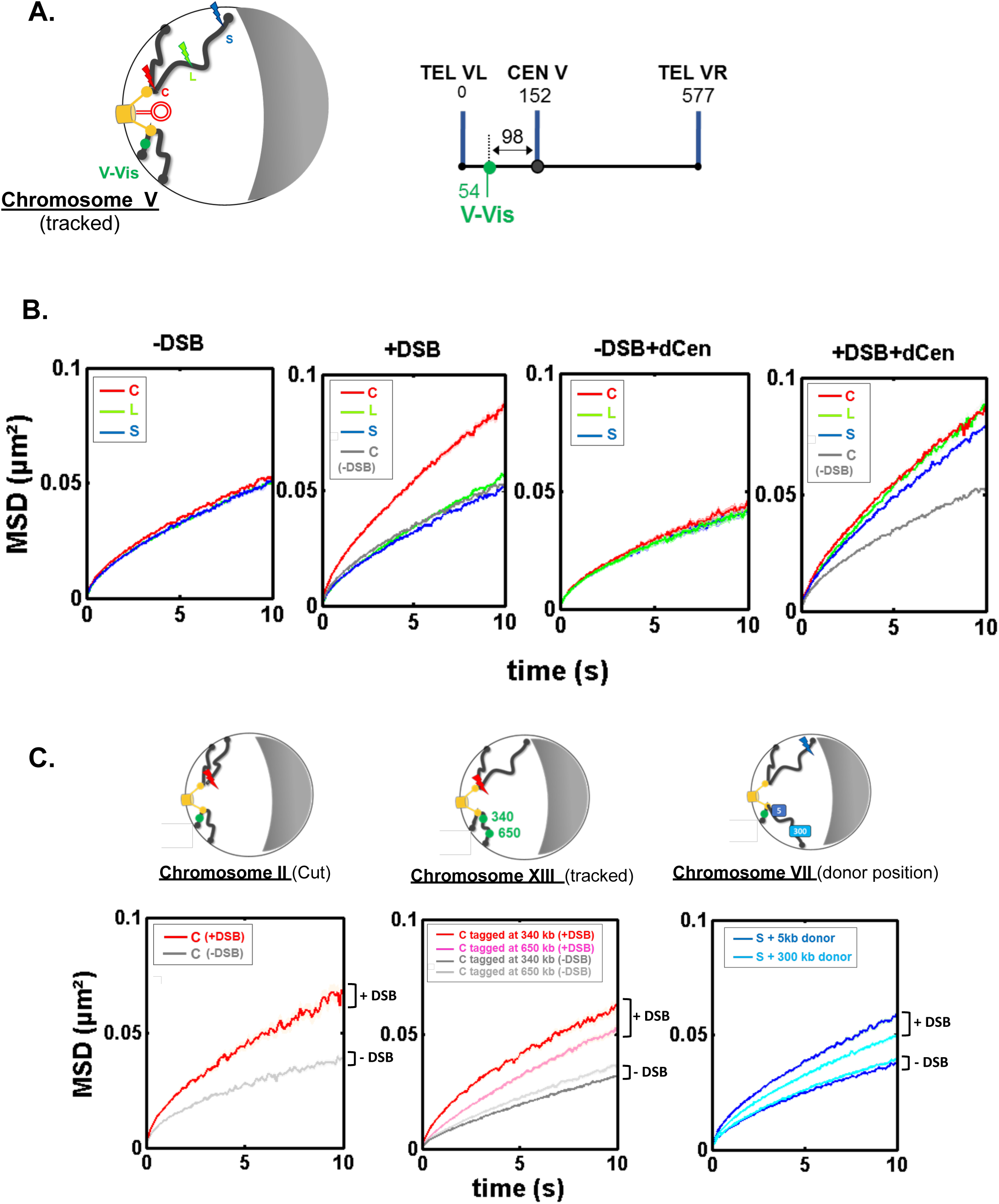
Enhanced global chromatin dynamics after damage is dictated by DSB and donor positions. **A**. As in Figure 1B, 2-D and linear schematics of the undamaged chromosome containing the V-Vis locus used for global mobility tracking, indicated as (tracked). V-Vis consists of TetO repeats bound by the TetR repressor protein fused to the fluorescent protein GFP, integrated into the *MAK10* locus on the left arm of chromosome V at 98kb from the centromere CENV (green dot). **B**. Mean squared displacements (MSDs) as a function of time for the V-Vis locus in strain C (red), L (green) and S (blue) respectively after 6 h in galactose medium in the absence or the presence of I-*Sce*I expressing plasmid (-DSB, +DSB) and in the presence of the donor plasmid (-DSB+dCen, +DSB+dCen plasmid), or the empty plasmid (-DSB, +DSB). MSDs are calculated from video microscopy data with a time sampling of Δt=100ms. The number of cells analyzed to calculate each curve is indicated in Table S1. The grey curve corresponds to the average MSD of the strain C without DSB in the presence of dCen or of the empty plasmid. **C**. The global increase in dynamics after DSB is verified for i) another pericentromeric DSB (left); ii) other tracked loci (middle) and iii) donor sequences distinct than dCen plasmid (right). i) I-*Sce*I cs is inserted at 8kb from CENII (red curve), MSD without DSB induction is shown in grey; ii) MSDs are calculated by tracking TetO repeats inserted on chromosome XIII at 340 and 650 kb of the centromere (red and pink curves, respectively) upon induction of I-*Sce*I DSB on peri-centromere of chromosome IV. The grey curves correspond to the average MSDs without induction for both strains; iii) the donor sequence is inserted on Chromosome VII (dChr.VII) after a DSB in subtelomeric position (S, blue lightening). MSDs are measured after insertion of donor sequence at 5kb or 300 kb of CEN VII (dark and light blue, respectively). Grey curves represent MSDs when DSB is not induced. 2-D schemes of Rabl chromosome conformation with positions of the DSB, the tracked locus or the donor sequences are shown on the top of each MSD graph as in Figure 1B.

Global mobility induced by a pericentromeric DSB, was not specific to Chromosome IV, since I-*Sce*I cs inserted at 7 kb from CENII induced a similar increase in MSDs of the V-Vis locus (**Fig. 2C, Chromosome II, cut**). Likewise, the increase in global mobility after a DSB in the C strain was observed elsewhere than at the V-Vis locus, on chromosome XIII at 340 and 650 kb from CENXIII, but to a lesser extent for the locus tracked away from the centromere (**Fig. 2C, Chromosome XIII, tracked**). Altogether these results show that in the absence of homology, the spatial position of the DSB plays a key role in induction of global mobility. A pericentromeric DSB specifically triggers a global mobility that reflects chromatin changes at short times and greater exploration of nuclear space at long times (Miné-Hattab et al. 2017). Furthermore, the gradual increase in mobility over the induction time suggests a progressive chromatin change behind global mobility at short times.

We then analyzed whether global mobility was affected during HR, by the presence of the dCen donor sequence. Before induction, dCen did not alter V-Vis mobility in any of the strains **(Fig. 2B, -DSB +dCen plasmid**). Upon DSB, in the presence of dCen, an increase of V-Vis MSDs was now observed even in L and S strains (**Fig. 2B** **and Supp. Fig 2C,+DSB +dCen plasmid**). We then asked whether mobility was also enhanced with an intra-chromosomal donor located on chromosome VII, near and far from the centromere (5kb and 300 kb from CENVII). Upon induction of the DSB in the S strain, the closer the donor was to the centromere, the greater was the V-Vis mobility (**Fig. 2C, Chromosome VII, donor position**). These observations highlight that in addition to chromosome architecture, sequence homology is instrumental in inducing global chromatin mobility, implying that global mobility may be triggered by homology search only when the DSB is distant from the pericentromeric domain.

### Enhanced chromatin mobility during Homologous Recombination is not dependent on cell cycle arrest but on DNA Damage signaling

If the increase in global mobility observed in the presence of a donor sequence could be explained by homology search, then the dynamic behavior of HR-repaired cells should be different from that of cells in search for homology. We therefore discriminated trajectories in the repaired and non-repaired cell populations. Whereas HR-repaired cells were red and cycling, white cells appeared blocked in G2/M, suggesting that these cells had not been repaired by NHEJ and were still searching for homology (**Fig. 3A**). MSD at 10sec. of V-Vis in white cells was higher than that of red cells (0.088 and 0.064, respectively), while cells without DSBs showed limited global mobility (MSD at 10sec. of 0.049) (**Fig. 3A** **and** **Fig. 3B**). To further determine whether the increased dynamics of unrepaired cells is due to their cell cycle arrest in G2/M, we measured V-Vis mobility in a *rad9Δ* checkpoint-deficient mutant. We first verified that I-*Sce*I cut efficiency was similar between WT and *Δrad9* strains and assessed cell survival and cell cycle progression in the *Δrad9* mutant (**Supp. Fig 3A**). As expected, *RAD9* deletion prevented cell cycle arrest and led to drastically altered cell growth after I-*Sce*I induction (**Supp. Fig 3B and Supp. Fig. 3C**).

**Figure 3.**
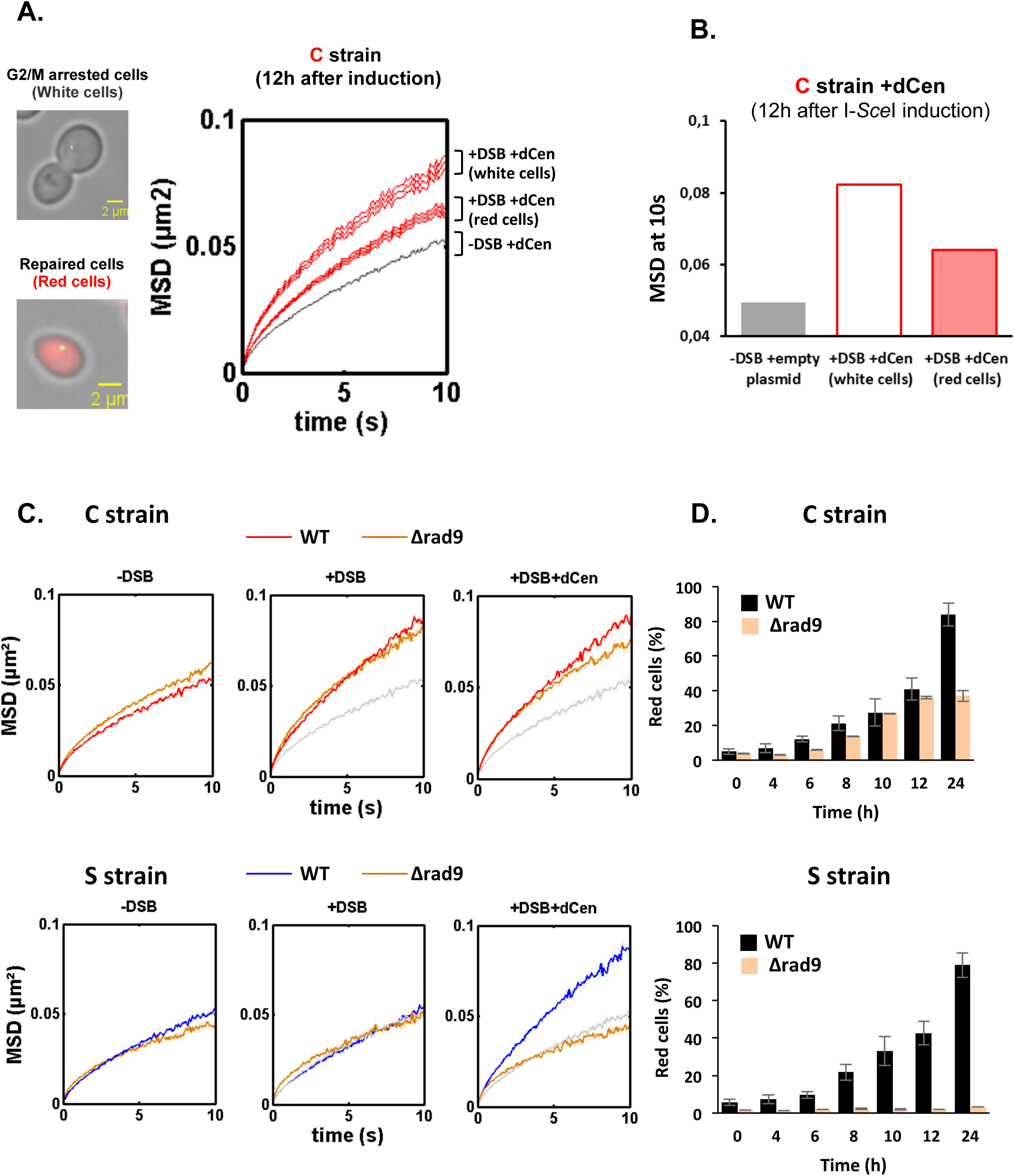
Proximal mobility is Rad9 independent. **A**. Non-repaired cells (white cells) show increased global mobility, while repaired cells (red cells) recover normal global mobility. MSDs of the V-Vis locus in strain C as a function of time, calculated as in Fig 2A. The grey and red curves correspond respectively to strain C in the presence of the donor dCen plasmid after 12 h in galactose medium with a plasmid not expressing I-*Sce*I (-DSB) and expressing I-SceI (+ DSB). MSDs are calculated after color visual cell sorting. Red MSD full curve corresponds to cycling red cells, red empty curve corresponds to white G2/M arrested cells. Examples of white, G2/M arrested and red, cycling cells are shown. Bar scale, 2µm. **B**. Box plots of MSDs at 10sec of undamaged, white and red cells 12h after I-*SceI* induction. The color code corresponds to that used in **A**. **C.** MSDs in strains C and S in WT and *Δrad9* mutant. MSDs are calculated as in Fig. 2 upon 6h of DSB induction with dCen plasmid or with empty plasmid (+DSB+dCen, +DSB respectively). Controls without DSB are also shown (-DSB). **D.** HR kinetics upon induction of I-*Sce*I in the presence of dCen in WT (black) and *Δrad9* (orange) C or S strains were measured by FACS as in Figure 1F. Error bars represent the standard deviation of three independent experiments.

When we measured global chromatin mobility upon DSB induction, we observed that surprisingly, its increase after pericentromeric DSB was similar between WT and *Δrad9* cells, regardless of whether dCen was present or not (**Fig 3C**). In contrast, no increase in global dynamics could be observed in the *Δrad9* mutant when DSB was induced away from the centromere, despite effective cleavage by I-*Sce*I (**Fig.3C, Supp. Fig 3A**). In the latter context, Rad9, therefore, became essential (**Supp. Fig 3B**), as observed when multiple damages were produced by Zeocin (Seeber et al. 2013; Garcia Fernandez et al. 2021). This indicates that the DNA Damage Response signaling is essential for global mobility only when long-range homology search is required.

Because the Rad9 defect prevents the cell cycle arrest required for repair, induction of a DSB in either the C or S strains resulted in cell death in the absence of a donor (**Supp. Fig.3B**). Furthermore, induction of a DSB resulted in cell death even in the presence of the donor sequence in the S strain (**Supp. Fig 3B**). In contrast, HR repair of the DSB generated in the pericentromeric region in the C strain was effective and 40% red cells were obtained after 24 h of induction (**Fig. 3D**).

These results exclude an obligatory effect of G2/M cell cycle arrest on both the global dynamics and HR repair. However, the requirement for Rad9 signaling is specific to the position of the DSB. Only the mechanism underlying pericentromeric proximal mobility could effectively proceed without Rad9.

### Enhanced global mobility is controlled by H2A phosphorylation

Phosphorylation of H2A occurs upstream of Rad9, is necessary and sufficient to induce global mobility when multiple damages occur and spreads through the centromeres after a pericentromeric DSB (Lee et al. 2014; Renkawitz et al. 2013b; Li et al. 2020; Garcia Fernandez et al. 2021). H2A phosphorylation was thus an attractive candidate to explain the mechanism by which global mobility occurred in a pericentromeric position-dependent manner. We therefore measured the distribution of ƴ-H2A(X) by chromatin immunoprecipitation (ChIP) in *cis* and in *trans* of the DSB before and after DSB induction, in C and S strains (**Fig. 4**). ChIP ƴ-H2A(X) signals in *cis* were detected by qPCR using a set of primers located at 5kb, 50kb and 100kb from the I-*Sce*I cutting site. ƴ-H2A(X) signal on the actin gene was used as a reference control. In the absence of DSB, low-intensity ƴ-H2A(X) ChIP signals were detected in either the C or S strains (**Fig.4A, grey bars**). Conversely, after 6h of I-*Sce*I induction, ChIP levels of ƴ-H2A(X) increased significantly in *cis* at 50 kb and 100 kb from the I-*Sce*I cutting site by approximately 15- and 10- fold, respectively, both in the C and S strains (**Fig. 4A, white bars**). No increase in ChIP ƴ-H2A(X) levels was detected at 5 kb of the cutting site, likely due to 5’ to 3’ resection occurring from the DSB ends (Eapen et al. 2012; Renkawitz et al. 2013b). In the presence of the donor, the enrichment of ƴ-H2A(X) was comparable to that observed with the empty plasmid, except at 100kb from the cutting site, where a decrease was observed, probably indicating repair (**Fig. 4A, red bars**). To next examine the levels of ƴ- H2A(X) relative to the centromere *in trans*, specific primers were designed at 15kb, 50kb and 100 kb in the region between the CENV and the V-Vis locus (**Fig. 4B**). In the absence of the donor, enrichment of ƴ-H2A(X) was observed only if DSB was induced near the centromere (C strain). ChIP levels of ƴ-H2A(X) increased six-fold in *trans* at 15kb to two-fold at 50kb and 100 kb from CEN-V (**Fig. 4B, white bars**). This is consistent with ƴ-H2A(X) propagation occurring across centromeres that are spatially close. Notably, the V-Vis locus, located at 90 kb from CENV, corresponds to a position where ƴ-H2A(X) ChIP signals were increased two-fold, suggesting a correlation between increased V-Vis mobility and higher ƴ- H2A(X) levels. Strikingly, an enrichment of ƴ-H2A(X) ChIP levels was found in *trans* with a similar increase in both C and S strains, in the presence of dCen, but not of the empty plasmid (**Fig. 4B**). Together, these results reveal that the presence of a donor sequence spatially close to the centromere is critical for the propagation of ƴ-H2A(X) in *trans,* independently of the genomic position of the DSB. They further support that ƴ-H2A(X) spreads around the break and across pericentromeric regions if the DSB is close to the centromere.

**Figure 4.**
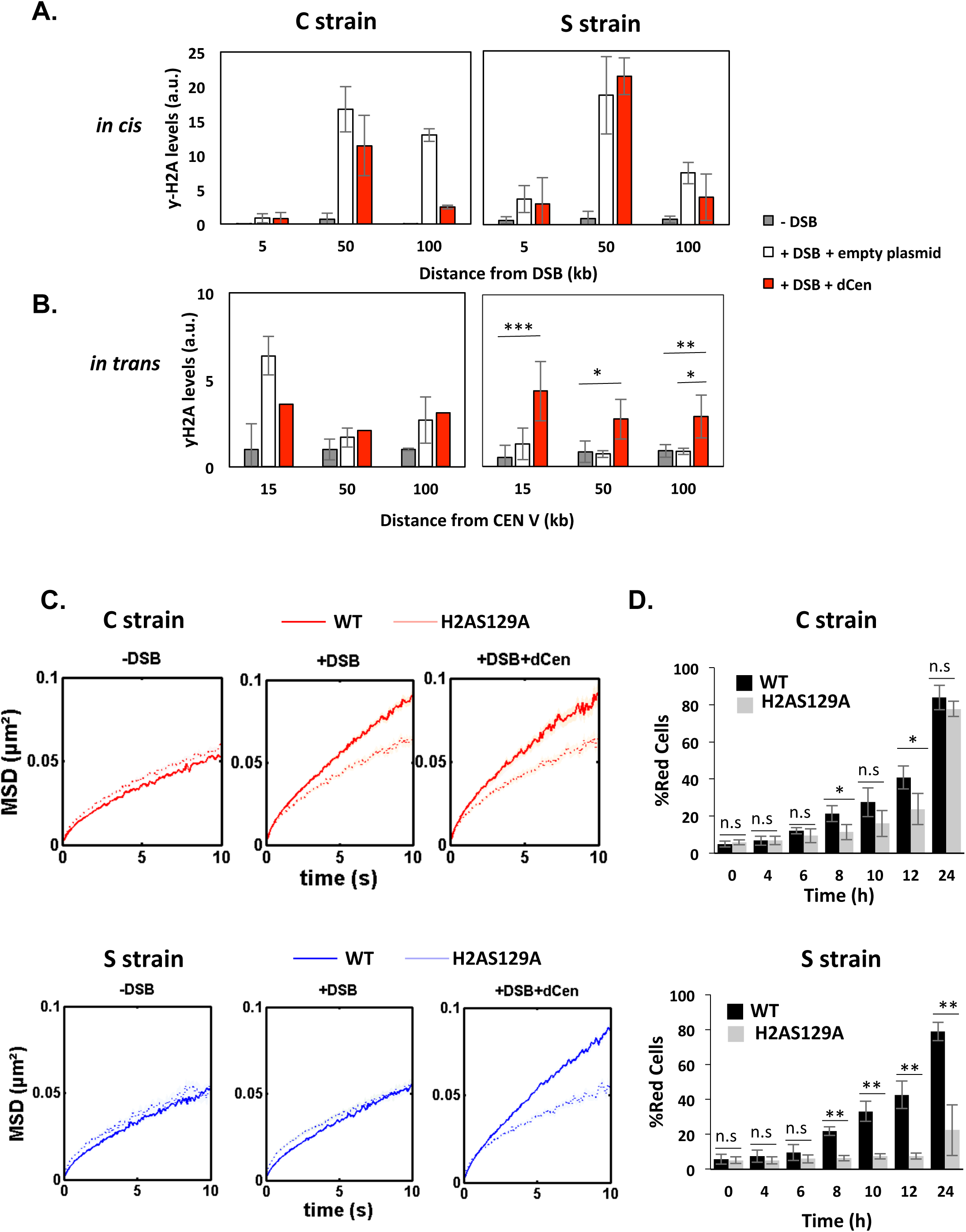
DSB induction in the presence of dCen triggers ƴ-H2A(X) spreading in trans. **A**. H2A phosphorylation spreading measured by ChIP around I-*Sce*I cs in strains C or S (*in cis*) in the absence (grey bars) or in the presence of DSB in the presence of the empty plasmid (white bars) or the dCen plasmid (red bars). ChIP was performed with ƴ-H2A(X) antibody after 6h induction of I-*Sce*I. Uncut control is shown as empty bars. DNA was analyzed by quantitative PCR using “*in cis*” primers corresponding to sequences at 5kb, 50kb and 100 kb from the right and left sides of I-*Sce*I cutting site in strains C and S, respectively. Actin was used as the reference gene for each condition. Each bar represents the ƴ-H2A(X) fold enrichment (ƴ-H2A(X)-IP/input relative to actin-IP/input) for undamaged and damaged conditions, respectively. The error bars represent the standard deviation of three independent experiments. **B**. ƴ-H2A(X) spreading “in *trans*” around pericentromeric region of chromosome V. ChIP was performed as above using “*in trans*” primers corresponding to sequences at 15kb, 50kb and 100kb from CENV. Note that V-Vis is positioned at 95.7kb from CENV. **C.** MSDs in strains C and S in H2A-S129A mutated backgrounds. MSDs are calculated as in Fig. 2 upon 6h of DSB induction with dCen or empty plasmid (+DSB+dCen, +DSB respectively). Controls without DSB are also shown (- DSB). Wilcoxon rank-sum test between distributions, with the p-value. n.s. non significant, * (p<0.05), ** (p<0.001). **D**. HR kinetics upon induction of I-*Sce*I in the presence of dCen in strains C or S in WT (black) and H2A-S129A (light grey) mutant were measured by FACS as in Fig. 1F. Error bars represent the standard deviation of three independent experiments. Wilcoxon rank-sum test between distributions, with the p-value. n.s. non significant, * (p<0.05).

Given the close correlation between H2A phosphorylation around V-Vis and the increased mobility at this locus, we further explored the involvement of ƴ-H2A(X) in mobility by measuring global dynamics in phospho-deficient H2A-S129A and phospho-mimetic H2A-S129E mutants, in the presence or absence of dCen. I-*Sce*I cutting efficiency in these mutated strains exhibited similar levels to the WT after 6h of I- *Sce*I expression, as shown by qPCR (**Supp. Fig. 4A**). Upon I-*Sce*I induction, serial dilutions of cells showed that C and S strains in the absence of dCen in either a wild-type or H2A-S129A mutant background were impaired in colonies formation (**Supp. Fig. 4B**). Yet, the H2A-S129E mutant showed better survival, as expected by its positive effect on NHEJ repair (Garcia Fernandez et al. 2021). In the presence of dCen, colonies formed in all strains, but were fewer in the S strain mutated for H2A-S129A, indicating that H2A phosphorylation is required for cell survival mainly when the DSB is distant from the centromere.

We next analyzed global mobility upon DSB induction in WT and H2A-S129 mutant strains. In the H2A- S129A mutant, the increase in global chromatin mobility was similarly impaired in the C and S strains harboring or not the donor sequence dCen (**Fig. 4C** and **Supp. Fig. 4E**). In contrast, the MSD curves of the phosphomimetic mutant H2A-S129E showed an increase in both C and S strains, even in the absence of DNA damage (**Supp. Fig. 4C** and **Supp. Fig. 4E**). These results clearly indicate that global mobility following a single DSB is primarily controlled by H2A(X) phosphorylation, presumably by spreading of this mark. This spreading may be triggered by a DSB near the pericentromeric domain, or by a centromeric donor when the DSB is induced elsewhere in the genome.

The kinetics of HR repair in H2A-S129 mutants was next assessed using the THRIV system (**Fig. 4D**). After a DSB generated near the centromere, the kinetics of red cells appearance was reduced in the H2A- S129A mutant compared with the WT, but the number of cells repaired by HR after 24h was comparable in both strains (**Fig. 4D, C**, **grey bars**). In contrast, in the same H2A-S129A mutant, a DSB generated in S strain could not be repaired by HR (**Fig. 4E, S, grey bars**). HR kinetics in the S129E mutant was similar to that in WT strains, regardless the DSB position (**Supp Fig. 4D**, **grey bars**). These results show that ƴ- H2A(X)-mediated mobility upon DSB induction is essential to facilitate HR. However, the need for mobility and 3D spatial sampling are determined by the position of the DSB and the donor sequence in the nucleus.

### Global chromatin mobility facilitates the search for a distant homologous donor within the nucleus

To further investigate whether global mobility was dependent on homology search, we tested the role of the repair protein Rad51 (Zou and Elledge 2003; Deng et al. 2015). It has been proposed that the 3’ single-stranded DNA extension covered by Rad51 probes homologies around the DSB and beyond within chromosomes, in a way that depends on their spatial organization (Renkawitz et al. 2013b). Rad51 has also been implicated in homology search in diploid cells (Mine-Hattab and Rothstein 2012; Miné-Hattab et al. 2017; Smith et al. 2018). It is further documented that ƴ-H2A(X) and Rad51 propagate concomitantly (Renkawitz et al. 2013b). Given our observations, Rad51 might be involved differently depending on the relative positions of the donor and recipient sequences. To test this, we deleted *RAD51* in the C and S strains, verified that cleavage by I-*Sce*I was as effective in both strains (**Fig. 5A**) and measured global dynamics in the presence or absence of damage (**Fig. 5B** and **Fig. 5C**). In the absence of a DSB, the global dynamics of WT and *Δrad51* strains were similar (**Fig. 5B** and **Fig. 5C, -DSB**). Upon I-*Sce*I induction, the *Δrad51* mutant strikingly did not affect the global mobility observed in the C strain (**Fig. 5B** and **Fig. 5C**, **C strain +DSB**). In contrast, deletion of *RAD51* abolished the increase in global dynamics induced in the S strain when dCen was present (**Fig. 5B** and **Fig. 5C**, **S strain +DSB +dCen**). These results substantiate that global mobility could require Rad51 for long-range homology search and highlight the existence of a Rad51-independent global mobility when the damage is close to the centromere.

**Figure 5.**
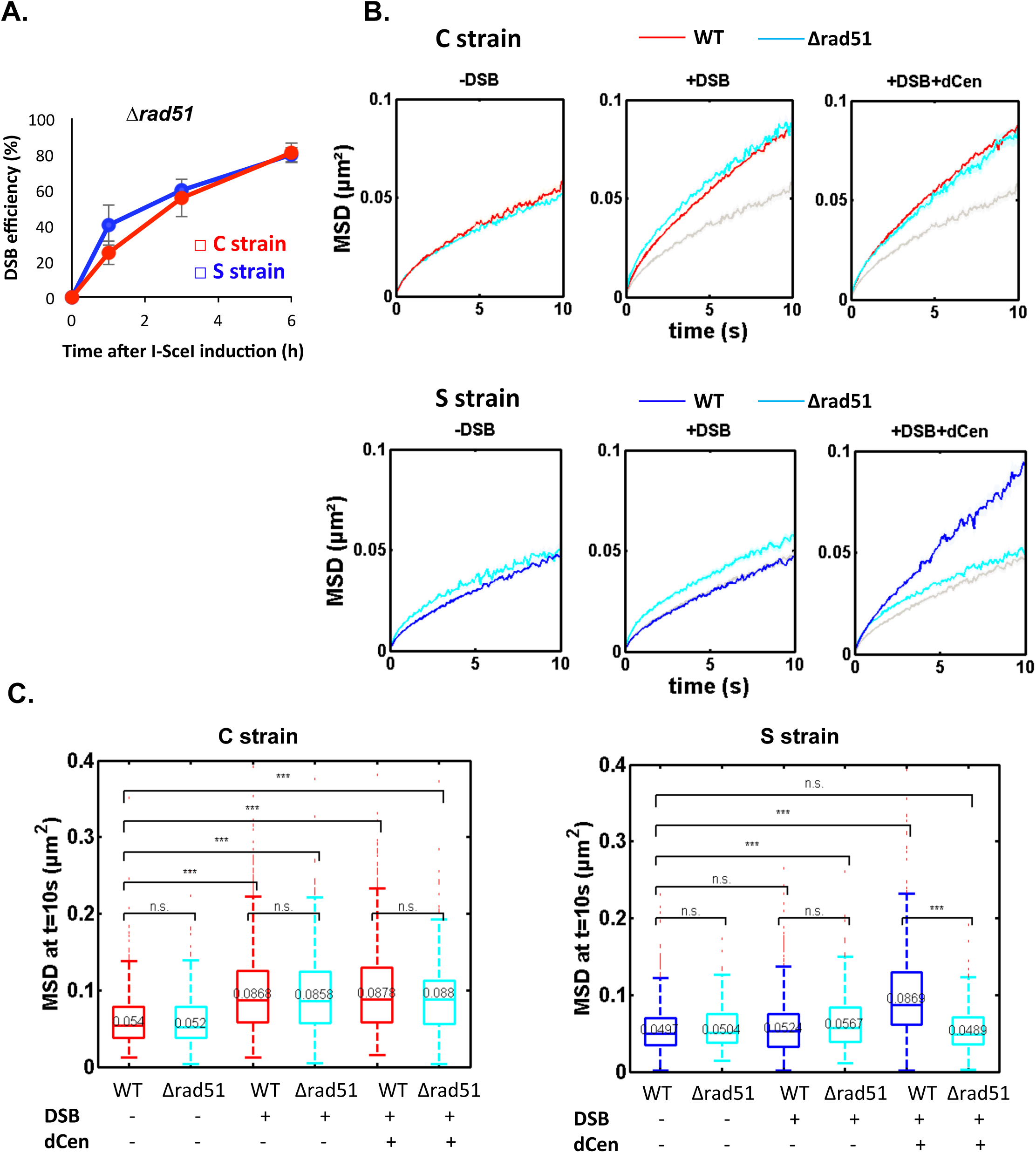
Global mobility requires Rad51 only when DSB and donor sequence are spatially distant. **A**. MSD of the V-Vis locus in strains C (red) or S (blue) mutated for Rad51 (*Δrad51*) after 6 h in galactose medium when in the presence of an empty plasmid, I-*Sce*I is not expressed (Left, - DSB), I-*Sce*I is expressed, (Middle, +DSB) or I-*Sce*I is expressed in the presence of dCen plasmid (Right, +DSB + dCen). The grey curve corresponds to the control without DSB (-DSB). I-*Sce*I cleavage efficiency is measured in *Δrad51* strains by qPCR using primers flanking the I-*Sce*I cutting site. The error bars represent the standard deviation of three independent experiments. **C.** Boxplots of the distribution of MSDs at 10s from Fig. 5A for strains C and S in WT and *Δrad51* background without or with a DSB (-DSB, +DSB), with dCen or empty plasmids (-dCen, +dCen). Median values, lower and upper quartiles are shown. Whiskers indicate the full range of measured values, except for outliers represented by small red dots. Parentheses indicate the result of a Wilcoxon rank-sum test between distributions, with the p-value. n.s. non significant, * (p<0.05), ** (p<0.001), *** (p<0.0001).

## Discussion

Increase of mobility of a genome damaged by DSBs is a universal response. How this occurs and what are the consequences for genomic integrity is not yet fully understood. In the presence of sequence homology, mobility could promote search for homologous sequences and help broken ends to navigate in the crowded nuclear space as proposed by (Smith et al. 2018; Cheblal et al. 2020; Renkawitz et al. 2013b), although chromatin mobility has been reported to be dispensable for recombination efficiency (Strecker et al. 2016). In yeast, the nuclear architecture of chromosomes corresponds to a Rabl configuration, where all centromeres anchored at one pole of the nucleus form a region where *trans* contacts are enriched (Duan et al. 2010; Berger et al. 2008; Lazar-Stefanita et al. 2017). The Rabl configuration has been described as critical for the efficiency of DSB repair. Indeed, sequences located in territories whose spatial proximity is defined by chromosome organization, were shown to be repaired by homologous recombination (HR) more efficiently than spatially distant regions (Agmon et al. 2013; Batté et al. 2017; Lee et al. 2015). However, it is not known if enhanced dynamics of the damaged or undamaged loci has a role in this repair efficiency. Here, we provide direct evidence that two types of global mobility are involved in HR in a chromosome organization-dependent manner.

Our results indicate that chromosome organization is critical in the establishment of global mobility. First, when a DSB is induced in a region where *trans* contacts are facilitated, such as near the centromeres, we observed an increase of mobility of other undamaged loci that we propose to name “proximal mobility”. This mobility in *trans* correlates with and depends solely on H2A phosphorylation that can spread from the site of damage to other pericentromeric regions. This was shown by ChIP in the strain where the break is induced near the centromere (strain C), in agreement with (Renkawitz et al. 2013b; Li et al. 2020). This proximal mobility is not altered by the presence of a donor sequence, or by checkpoint progression, as shown by the lack of effect of *RAD9* deletion (53BP1 ortholog) on mobility. This indicates that at least under these conditions, the ƴ-H2A(X) spreading induced by nuclear organization is sufficient to enhance chromatin mobility. ƴ-H2A(X) spreading *in trans* within the pericentromeric region is mainly induced by the H2A kinase Mec1 (Renkawitz et al. 2013b; Lee et al. 2014; Li et al. 2020). A recent study combining experimental ƴ-H2A(X) ChIP in *Δmec1* mutant and mathematical modeling provides a new perspective of the way Mec1 may diffuse from the site of damage by diffusion in 3D space (Li et al. 2020). Our observation that proximal mobility is only dependent on ƴ-H2A(X) is consistent with this model of Mec1 diffusion across centromeres. The attenuated Mec1 diffusion along the chromatin fiber reported by Li et al, could also explain why tracking loci further away from the centromere, at 340kb or 650kb from it, reveals a smaller increase in global mobility (**Fig. 2C, Table S1**). It would be interesting to test, with polymer models, how this ƴ-H2A(X)- induced chromatin modification allows the propagation of movement to chromosomal regions further away from the centromere.

In addition to enriched *trans*-contacts, the global mobility effect observed near the centromere could be related to a specific chromosome organization in this region. Indeed, centromeres are enriched in structural maintenance of chromosomes (SMCs) complexes, including cohesins and Smc5/6, with a strong impact on chromatin loop formation and size (Dauban et al. 2020; Piazza et al. 2021; Weber et al. 2004; Betts Lindroos et al. 2006; Lazar-Stefanita et al. 2017; Paldi et al. 2020). Thus, cohesins, also recruited during DSB repair (Ünal et al. 2007; Ström et al. 2004; Betts Lindroos et al. 2006), could promote H2A phosphorylation during loop extrusion, creating a functional repair unit, as it has been proposed for mammalian TADs (Arnould et al. 2021). In this latter study, the Tel1 ortholog ATM, was proposed to remain attached to the broken ends of the DNA, promoting progressive and directional phosphorylation of nucleosomes compatible with a loop mechanism of DNA extrusion (Arnould et al. 2021). If the loop model may not match with the bending properties of yeast chromatin (Li et al. 2020), the gradual increase in mobility we observed in the strain C is consistent with progressive phosphorylation of H2A along the chromosome over time. The pericentromeric environment beneficial to global mobility could also arise from microtubule-centromere links, although loss of centromeric tethering as a source of global mobility is still debated (Cheblal et al. 2020; Strecker et al. 2016; Lawrimore et al. 2017). The pericentromeric domain could reflect the 20-50kb region flanking the conserved 125bp of the centromere, where cohesins are preferentially enriched (Lazar-Stefanita et al. 2017; Dauban et al. 2020; Paldi et al. 2020). The specificity of pericentromeric chromatin could thus be explained by the characteristic Rabl configuration in yeast, where nuclear microtubules emanating from the yeast MTOC, associate with the kinetochores, required for cohesin association (Weber et al. 2004). The recent discovery that Rabl-like configurations emerged repeatedly throughout evolution, further suggests a functional advantage to centromeric clustering (Hoencamp et al. 2021).

We describe a second type of global mobility, which we propose to call “distal mobility”, when the damage occurred spatially far from the donor sequence, such as at the middle or end of the IVR chromosome arm (strains L and S). In this case, the increased mobility of V-vis is induced only if a donor close to the centromere, either on a plasmid or on the chromosome, is present. In this situation, a ƴ- H2A(X) enrichment is surprisingly observed in the vicinity of V-Vis. Strikingly, this donor-dependent distal mobility requires a more sophisticated mechanism than simple ƴ-H2A(X) propagation, since it involves checkpoint progression and is triggered in a Rad51-dependent manner. The requirement of Rad51 suggests that the nucleofilament and global mobility together are involved in homology search, in agreement with what was found by Rad51 ChIP (Renkawitz et al. 2013b). How does distal mobility related to homology search proceed? We know from ChIP of ƴ-H2A(X) and from H2A-S129 mutants that γH2A(X) signal spreading and homology search are apparently directly linked (**Fig. 4** and Renkawitz et al., 2013). The fact that the presence of a donor is required to observe both H2A phosphorylation and increased mobility seems to confirm the sampling of the genome by the Rad51 nucleofilament, together with the Mec1 as proposed by (Renkawitz et al. 2013b, 2013a). Furthermore, it implies the recombination process itself in ƴ-H2A(X) spreading near the donor. Phosphorylation could promote mobility through a stiffer chromatin, due to the repulsion of phosphate negative charges, as we have proposed (Herbert et al. 2017; Garcia Fernandez et al. 2021). H2A phosphorylation could also differentially modify chromatin structure by modulating the recruitment and/or the stable docking of DNA-binding proteins, such as INO80, a chromatin remodeler complex important in nucleosome stability (Cheblal et al. 2020; Hauer et al. 2017; Van Attikum et al. 2004; Bennett et al. 2013), or SMC complexes that could regulate loop formation and hence chromatin structure and mobility (Mirny and Dekker 2021; Cheblal et al. 2020). Recently, Piazza et al, found that a DSB involves cohesin-dependent scaffolding of the broken chromosome, which isolates this chromosome from others and inhibits *trans* contacts (Piazza et al. 2021). The distal mobility we describe here could help overcome this insulation and facilitate *trans* homology search when needed.

In addition to being induced by distinct but not exclusive mechanisms, the proximal and distal mobility described here have different roles in HR repair, as documented by directly tracking cells repaired by HR *in vivo*. While distal mobility is linked to HR repair, proximal mobility serves as a dispensable booster (**Fig. 6**). Indeed, H2A-S129A mutants are able to form a functional mCherry by HR, and therefore red colonies, when the cut is centromere-proximal (C strain), although proximal mobility is not increased. Red cells appear more slowly than in the wild type but reach a comparable level between the two strains after 24 hours. Thus, HR can take place without the genome being more mobile, but proximal mobility accelerates it. Furthermore, in the strain C deleted for *RAD9* checkpoint gene, red cells surprisingly appear, although not as abundant as in WT strain. In *Δrad9* mutant, proximal mobility is not affected but cell cycle arrest required for efficient HR is inefficient, suggesting that proximal mobility suppresses the recombination defects associated with the absence of Rad9, at least partially. This observation is remarkably consistent with our recent findings establishing that the global mobility observed in the H2A-S129E mutant suppresses the *Δrad9* checkpoint defect (Garcia Fernandez et al. 2021). The Rad9 checkpoint appears to play an ambivalent role in global mobility that depends on the nature of the damage and the mode of repair (Seeber et al. 2013; Garcia Fernandez et al. 2021). In situations where repair is challenging, such as spatially distant donor and recipient sequences (such as in the strain S) or when the cell experiences multiple random damages, the Rad9 checkpoint will collaborate for global chromatin dynamics as part of damage signaling. It will be essential to understand by which mechanism Rad9 defect is suppressed when chromatin becomes more mobile.

**Figure 6.**
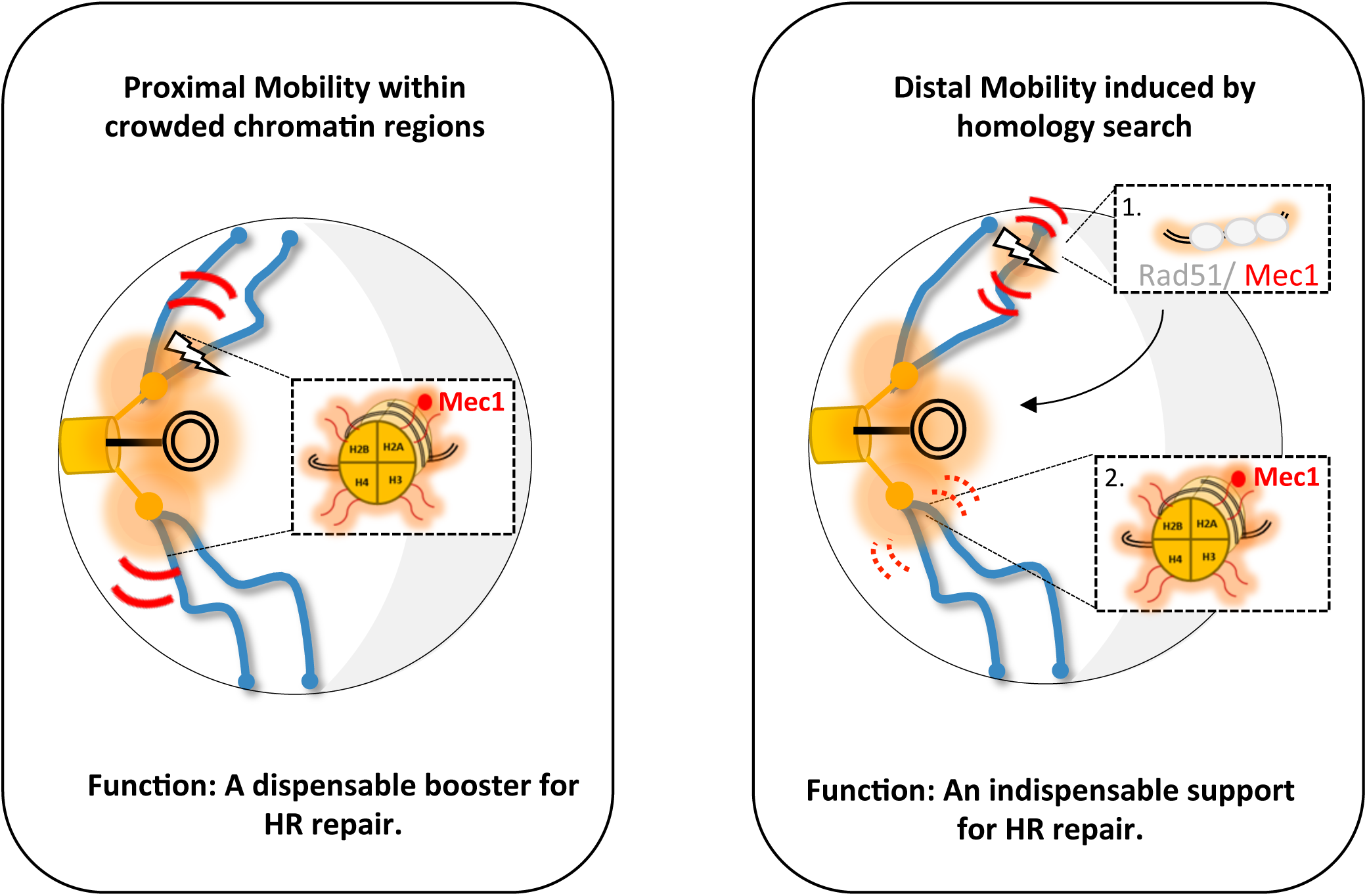
Two types of global mobility are involved in DSB repair by homologous recombination. A proximal mobility (left), happens in nuclear domains where *trans*-contacts are enriched, as in the pericentromeric region. Only the phosphorylation of H2A is involved in this global mobility, which serves to boost the speed of HR. Distal mobility (right) happens when the DSB is initiated in a region far from the centromeres. The Rad51 nucleofilament is required, as well as the HR to promote the phosphorylation of histone H2A. The ƴ-H2A(X) embedded chromatin is represented by the orange shadow, at the break site and in *trans* in the pericentromeric region. Red curves represent mobility whose thickness symbolizes the movement amplitude, fine curves for global mobility and thick curves for local mobility. The lightning bolt represents the double strand break.

Chromosome organization influences chromatin mobility when damage occurs, and this can accelerate repair either by NHEJ (Ma et al. 2021; Garcia Fernandez et al. 2021) or by HR as shown here. The global chromatin mobility observed in different regions of the nucleus could thus be a way to coordinate different responses to DNA damage. The proximal mobility observed in the pericentromeric region, in particular, may be an additional means of ensuring the essential maintenance of centromeres (Yilmaz et al. 2021). Interestingly, a sophisticated biological function such as sleep has been associated with chromosome mobility (Zada et al. 2019). In zebrafish, a link was found between neuronal activity, chromosome mobility and the number of lesions, also highlighting the protective role of chromosome mobility. Universally, global mobility would be an additional warrant of genomic integrity.

**Table S1** Compilation of values extracted from Mean squared displacements of the analyzed strains

**Table S2** Genotype of strains used in this study

## Materials and Methods

### Yeast culture

Yeast cells were grown in rich medium (yeast extract-peptone-dextrose, YPD) or synthetic complete (SC) medium lacking that appropriate amino acid at 30°C. Synthetic medium containing 2% raffinose and lacking appropriate amino acids was used to grow the cells overnight prior the induction of I-*Sce*I by plating onto 2% galactose plates or addition of 2% galactose in liquid culture. For high through put *in vivo* spot tracking, cells were plated on agarose patches (made of synthetic complete medium containing 2% agarose and galactose) and sealed using VaLaP (1/3 Vaseline, 1/3 Lanoline and 1/3 Paraffin).

### Yeast Strains and Plasmids

Strains C, L, and S were constructed by first integrating 256 TetO sequences at the MAK10 gene (YEL053C) as in (Strecker et al. 2016). To insert I-*Sce*I cutting site at the desired position, a *KANMX* cassette was integrated into intergenic loci of chromosome IV at positions 453880 + 453959 (C strain); 853996 + 854075 (L strain); 1511840 + 1511919 (S strain). The *KANMX* cassette was then replaced by homologous recombination using CRISPR/Cas9 (pEF526) with the amplified donor sequence mCherry* containing cutting site of I-*Sce*I (5‘-ttacgctagggataacagggtaatatagcg-3 ‘), from pCM189-mCherry*-I-*Sce*I (cs), with oligos oEA310/311. The dCen (pEF573) plasmid was obtained by PCR amplifying mCherry sequence with (oEA307/308) oligonucleotides and cloned into pRS412 (*ADE2* centromeric plasmid) digested with *BamH*I and *EcoR*I. pB07 plasmid containing I-*Sce*I under the control of a *GAL1-10* promoter in a *HIS3* plasmid was obtained from B. Pardo.

The H2A-S129A and H2A-S129E mutants were constructed as described in (Garcia Fernandez et al. 2021). Briefly, the CRISPR/Cas9 containing plasmids (pEF567 and pEF568) including the guide RNA (gRNA) targeting *HTA1* (oEA325/326) and *HTA2* (oEA327/328) inserted into the plasmid pJH2971 (CRISPR/Cas9- KANMX-Addgene plasmid#100955) were used to generate a DNA break, afterwards repaired with a donor oligonucleotide of 80nt carrying the desired mutation. All integrations were confirmed by PCR and mutations checked by sequencing. The primers and RNA guides used are indicated in Table S2.

Deletions of *RAD9* (YDR217C) and *RAD51* (YER095W) genes in the different strains were carried out by integration of the amplified *KANMX* cassette derived from the Saccharomyces Genome Deletion Project collection.

### Drop assays

For drop assays, overnight cultures were diluted to a starting density of OD600 = 1.0 and serial 1:5 dilutions were plated on selective medium (SC-Ade-His) containing raffinose or galactose (2%), and incubated at 30°C for 36h.

### Microscopy wide-field conditions

Live cell imaging was done using a widefield microscopy system featuring a Nikon Ti-E body equipped with the Perfect Focus System and a 60× oil immersion objective with a numerical aperture of 1.4 (Nikon, Plan APO). We used an Andor Neo sCMOS camera, which features a large field of viewof 276 × 233 μm at a pixel size of 108 nm. We acquired 3min films consecutively with an exposure time of 100 ms. The complete imaging system including camera, piezo, LEDs (SpectraX) was controlled by the NIS element software (version 4.60).

### Image analysis and statistics

Particle tracking and MSD analyses were performed as described in (Garcia Fernandez et al. 2021). Briefly, a spot-tracking algorithm (Fiji macro) allows isolating each spot and extracting its trajectory from the 2-D time-lapse sequences. MATLAB scripts are then applied to correct global displacements, control signal-noise ratio and compute MSD (Mean Squared Displacements) curves for each trajectory using non-overlapping time intervals and power laws fitting to population-averaged MSD over 10s intervals

### qPCR and RT-qPCR

Cells were grown to exponential phase in Raffinose selective medium and induced by addition of Galactose 2% final concentration. 1 to 2.10^8^ cells were centrifuged for 1 min at 13000 rpm and 300mg of glass micro beads, 200 μL of lysis solution (2% Triton X-100, 1% SDS, 100 mM NaCl, 10 mM Tris pH 8, 1 mM EDTA) and 200 μl of phenol-chloroform-isoamyl alcohol were added to the cell pellet. The mixture was vortexed for 10 min. at 4° C and centrifuged for 10 min. at 13,000 rpm. 140 μL of the aqueous phase were transferred into new tubes containing 500 μL of cold 100% ethanol. The mixture was centrifuged for 10 min. at 13,000 rpm at 4° C, the pellet washed with 500 μL of 70% ethanol, dried at room temperature, resuspended in 40 μL of TE containing 20 μg/ml of RNAse and incubated at 42 ° C for 30 min. The DNA was quantified by SimpliNano^TM^ fromBiochrom.

For RT-qPCR, RNA was extracted with the Machery Nagel kit, quantified by Nanoview and absence of RNA degradation checked by gel electrophoresis migration. Maxima First Strand was used for RT reaction on 1.5µg RNA.

The qPCR was performed on LightCycler® 480 (Roche) on a 96-well plate under the following conditions: Pre-incubation (95°C - 5 min. - 4,4 °C/min), amplification (45 cycles; 95°C – 10 sec. - 4,4 °C/min.; 58°C – 10 sec. - 2,2 °C/min.; 72°C – 20 sec. - 4,4 °C/min.), melting curve (95°C – 5 sec. - 4,4 °C/min.; 60°C – 1 min. - 2,2 °C/min.; 97°C – continuous - 0,06 °C/min; 10 acquisitons/°C) and cooling (40°C – 30 sec. - 2,2 °C/min.).

The primers oEA312 / 313 (table S2) were used for qPCR and the actin gene as amplification control.

### Chromatin Immunoprecipitation

This protocol is adapted from (Forey et al. 2020), 1×10^9^ cells were cross-linked for 10 min with 1% formaldehyde (Sigma F8775) at room temperature on a shaking device. Fixation was quenched by addition of 0.25 M glycine (Sigma G8898) for 5 min. under agitation. Cells were washed two times with cold TBS1X (4°C). Dry pellets were frozen and stored at -20°C. Cell pellets were resuspended in lysis buffer (50 mM HEPES-KOH pH7.5, 140 mM NaCl, 1 mM EDTA, 1% Triton-X100, 0.1% sodium deoxycholate) supplemented with 1 mM PMSF, phosphatase inhibitor and anti-protease (complete Tablet, Roche, 505649001) and lysed three times by beads shaking. The lysate (WCE, Whole Cell Extract) volume was sonicated with a Q500 sonicator (Qsonica; 3 cycles: 40 sec ON, 40 sec OFF, amplitude medium). 20 μL of input material were saved for qPCR. Approximately 180μL (13 OD) of the input were incubated with 0.5% BSA, 1μL of DNA carrier (10mg/ml) and 2 μL of anti-γ-H2A(X) (Abcam 15083) on a rotating wheel overnight at 4°C. The day after, 30 μL of protein G Sepharose beads were washed three times and resuspended in 90 μL final of lysis buffer, which were added to the overnight culture during 2h on a rotating wheel at 4°C. Beads were then collected and washed on ice with cold solutions: twice with Lysis buffer (50 mM HEPES-KOH pH7.5, 140mM NaCl, 1mM EDTA, 1% Triton115 X100, 0.1% sodium deoxycholate), twice with Lysis buffer added with 360mM NaCl, twice with Washing buffer (10 mM Tris- HCl pH8, 0.25 M LiCl, 0.5% IGEPAL, 1mM EDTA, 0.1% sodium deoxycholate) and once with TE (10mM Tris-HCl pH8, 1mM EDTA). Antibodies were uncoupled from beads with 150 μL of Elution Buffer (50mM Tris-HCl pH8, 10mM EDTA, 1% SDS) for 20 min at 65°C. Eluates were incubated with 120 μL of TE containing 0.1% SDS at 65°C for 6 hours to de-crosslink. Then 130 μL of TE containing 60 μg RNase A (Sigma, R65-13) were added and the samples were incubated for 2 hours at 37°C. Proteins were digested by addition of 20 μL of proteinase K (Sigma, P6556) at 20 mg/ml to the samples and incubation for 2 hours at 37°C. DNA purification was completed with the QIAquick® PCR Purification Kit (n28104). Finally, qPCR reactions were performed in a LightCycler480 (Roche). IP/Input ratios were calculated and qPCR results were normalized on ChIP-qPCR Act1 for ƴ-H2A(X). For each strain, three independent experiments were performed with the corresponding controls.

### FACS

For cell cycle, cells were grown to mid-log-phase in liquid cultures, and treated or not with Zeocin at 250 mg/μL during 6h at 30°C. After incubation, samples were fixed with 70% ethanol and kept at 4°C for 48 h. Cells were then resuspended in 50 mM Sodium Citrate (pH 7) containing RnaseA at 0.2 mg/mL final concentration. After incubation at 37°C for 1h, Sytox Green was added to a final concentration of 1mM. A total of 10^6^ cells were analyzed with a CANTO II flow cytometer (BD Biosciences). Aggregates and dead cells were gated out, and percentages of cells with 1C and 2C DNA content were calculated using FLOWJO software.

For the kinetics of fluorescence red cells, 1 to 2.10^7^ cells were removed and centrifuged. All the following steps were done in the dark. The pellet was washed once with cold PBS at 4° C. After removal of the supernatant, the pellet was resuspended in 500 μL of cold PBS and 500 μL of cold PBS with formaldehyde (to a final concentration of 1%) were added and gently mixed. The cells were incubated for 15 min at 4° C. The cells were then washed 3 times with sodium citrate (50 mM) and prepared at a concentration of 1.10^6^ cells / ml for the cytometry carried out immediately after (in order to limit the loss of fluorescence). Following steps were performed as above.

## Acknowledgments

We thank Benjamin Pardo for his advices in setting up the chromatin immunoprecipitation protocol and Stanford and Sup BioTech internships Tara Shanon, Dahee Chung and Noémie Guerre for their participation in different constructs steps of this study. We acknowledge the critical and constructive reading of Amandine Bonnet, Judith Miné-Hattab and Pascale Lesage. We thank Jean Michel Arbona and the team members for numerous discussions.

## Fundings

This research was funded by the Agence Nationale pour la Recherche, (ANR-17-CE11-0025 to E.F.), the initiatives d’excellence (Idex ANR11-LABX-0071, IDEX-0005-), the Institut National du Cancer (INCA), grant number PLBIOR21018HH and Fondation ARC pour la recherche sur le Cancer, grants number PJA32020060002313 and DOC20190508798. F.G.F. acknowledges the Peruvian Scholarship Cienciactiva of Consejo Nacional de Ciencia, Tecnologıa e Innovación Tecnológica (CONCYTEC) for supporting her PhD study at INSERM and Diderot University and support by Fondation ARC pour la Recherche sur le Cancer (DOC20190508798). E.A. acknowledges the IDEX USPC (NUPGE15RDX), A.C.S and Y.K. the Fondation pour la Recherche Médicale FRM (ECO202006011576 and ING20160435205 respectively).

## Author contributions

FGF, EA, ACS, RB ,YK and MB performed the experiments; FGF, EA, ACS, RB , YK, MB and EF analyzed and interpreted the data; FGF, EA and EF conceive and designed the work; FGF and EF wrote the paper; EF secured funding.

## Conflict of interest

Authors declare no conflict of interest

---

**Supplementary Figure 1.**
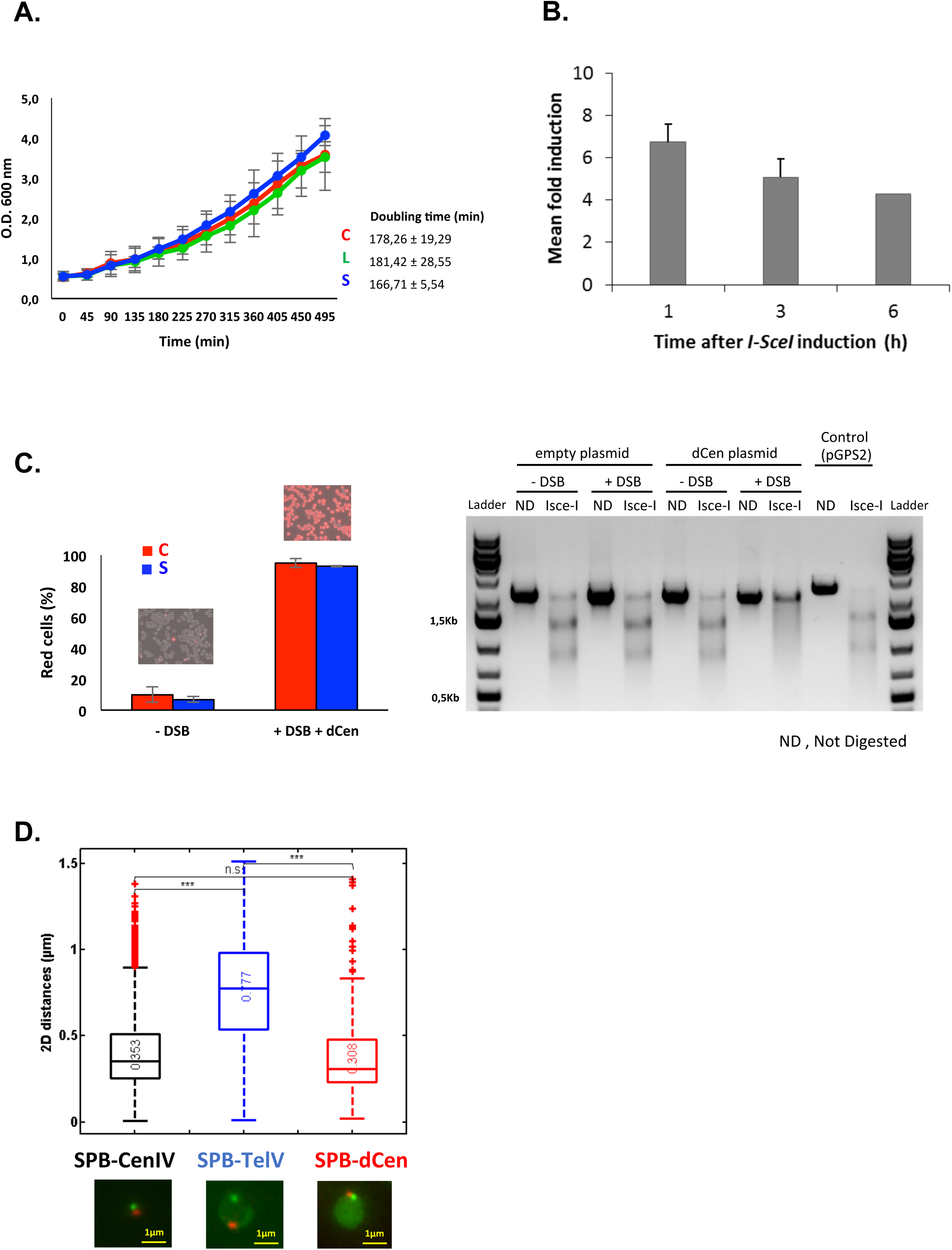
**A**. Growth curve of strains C (red), L (green) and S (blue) showing a similar doubling time. **B**. I-*Sce*I induction ratio measured by Transcription Quantitative Reverse PCR (RT-qPCR) using primers hybridizing into the I-*Sce*I coding sequence. The error bars represent the standard deviation of three independent experiments. **C**. Quantification of red and white cells in strains C and S, 48h after I-*Sce*I induction on galactose plates before DNA extraction and PCR amplification around the I-*Sce*I cs of those strains. If the recipient I-*Sce*I cs containing sequence is fully repaired by homologous recombination with the donor dCen plasmid, it cannot be cut by I-*Sce*I *in vitro*. pGPS2 I-*Sce*I cs containing plasmid is used as a control for *in vitro* digestion. **D**. Boxplot of 2D distances measured in μm between the SPB (SPC29-mCherry, red) and dCen plasmid containing tetO repeats, labeled in green (TetR-GFP). As references, chromosome loci labeled either close to the centromere (YDR003w, 5kb from CenIV), or the telomere (YER188C, 8kb from TELVR) are shown. Below, examples of cell nuclei with the two red and green labeled loci allowing calculation of the SPB-loci distances. As expected, the donor sequence dCen on the centromeric plasmid is located close to the SPB, the subtelomeric sequence far from it, see (Therizols et al. 2010).

**Supplementary Figure 2.**
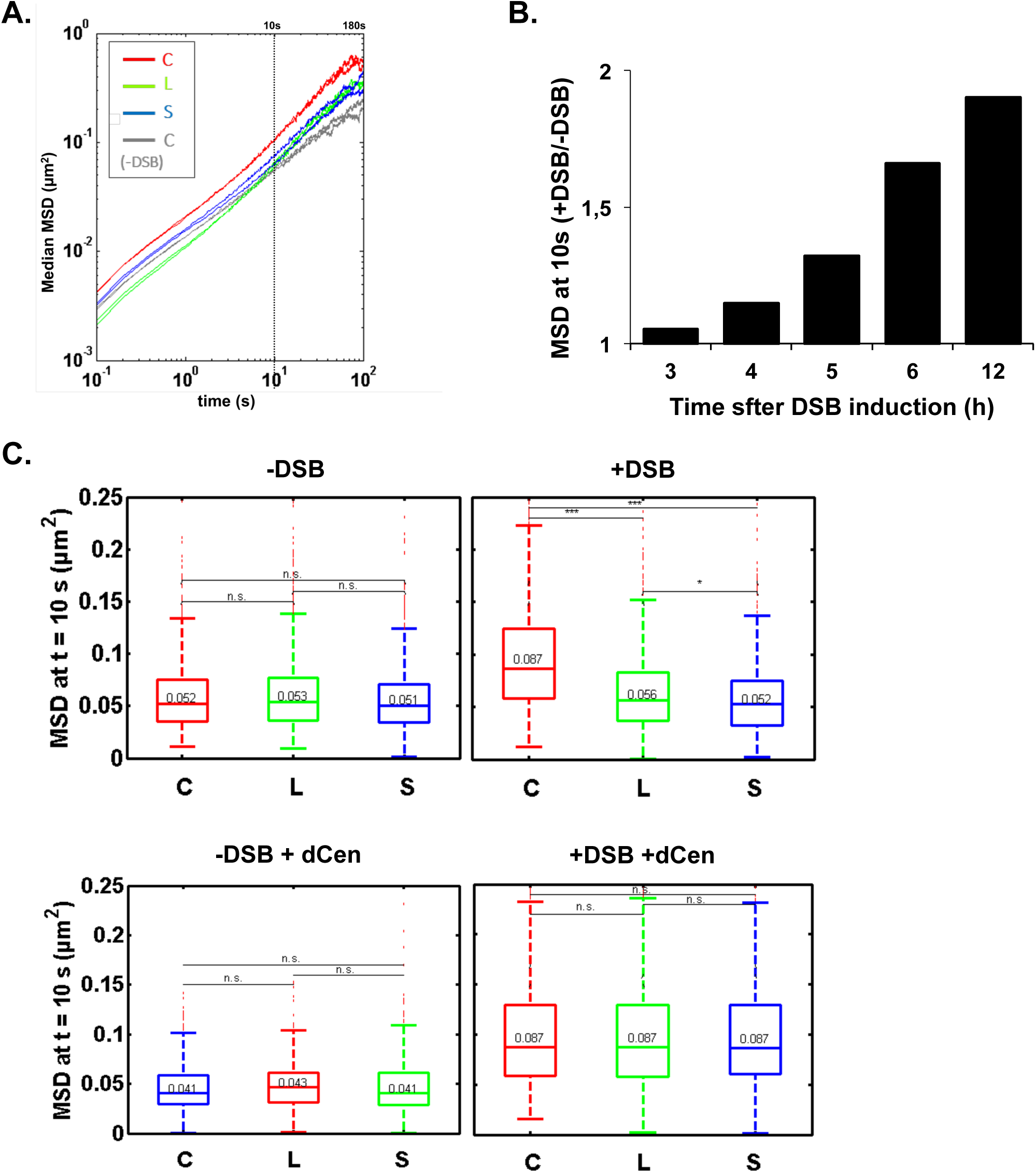
**A**. MSDs of strains C (red), L (green) and S (blue) respectively in the absence of donor sequence at long time points. **B**. Gradual increase of MSD with induction time in the strain C in the absence of donor. The relative ratio of MSDs at 10sec (+DSB/-DSB) for each time point after induction of I-SceI are shown. **C**. Boxplots of the distribution of MSDs at 10s from figure 2B for strains C, L and S with the same color code. Median values, lower and upper quartiles are shown. Whiskers indicate the full range of measured values, except for outliers represented by small red dots. Parentheses indicate the result of a Wilcoxon rank-sum test between distributions, with the p-value. n.s. non significant, * (p<0.05), ** (p<0.001), *** (p<0.0001).

**Supplementary Figure 3.**
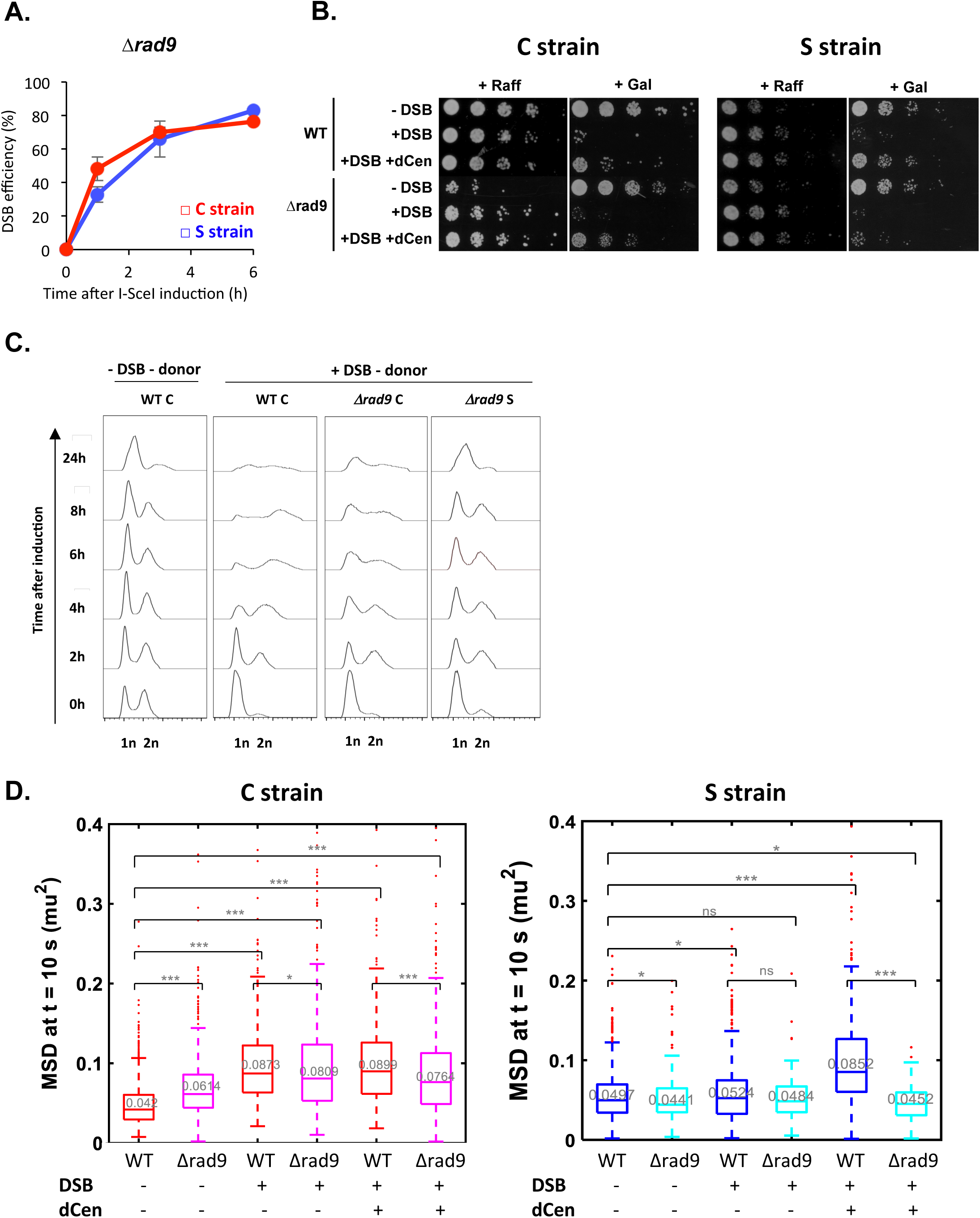
**A**. I-*Sce*I cleavage efficiency measured in *Δrad9* mutant by qPCR using primers flanking the I-*Sce*I cs. The error bars represent the standard deviation of three independent experiments. **B**. Drop assay (tenfold serial dilutions) showing the sensitivity of WT and *Δrad9* C or S strains, containing a plasmid expressing or not I-*Sce*I (-DSB, +DSB respectively), with dCen or an empty plasmid, when I-*Sce*I is induced (Galactose, Gal) or not induced (Raffinose, Raff). **C.** Analysis of the cell cycle of *Δrad9* C and S strains after induction of I-*Sce*I with dCen. Quantification is done as in Fig. 1.D. The peaks corresponding to 1n and 2n amount of DNA are indicated. **D.** Boxplots of the distribution of MSDs at 10s from Fig. 3C for WT and *Δrad9* C and S strains. Median values, lower and upper quartiles are shown. Whiskers indicate the full range of measured values, except for outliers represented by small red dots. Parentheses indicate the result of a Wilcoxon rank-sum test between distributions, with the p- value. n.s. non significant, * (p<0.05), ** (p<0.001), *** (p<0.0001).

**Supplementary Figure 4.**
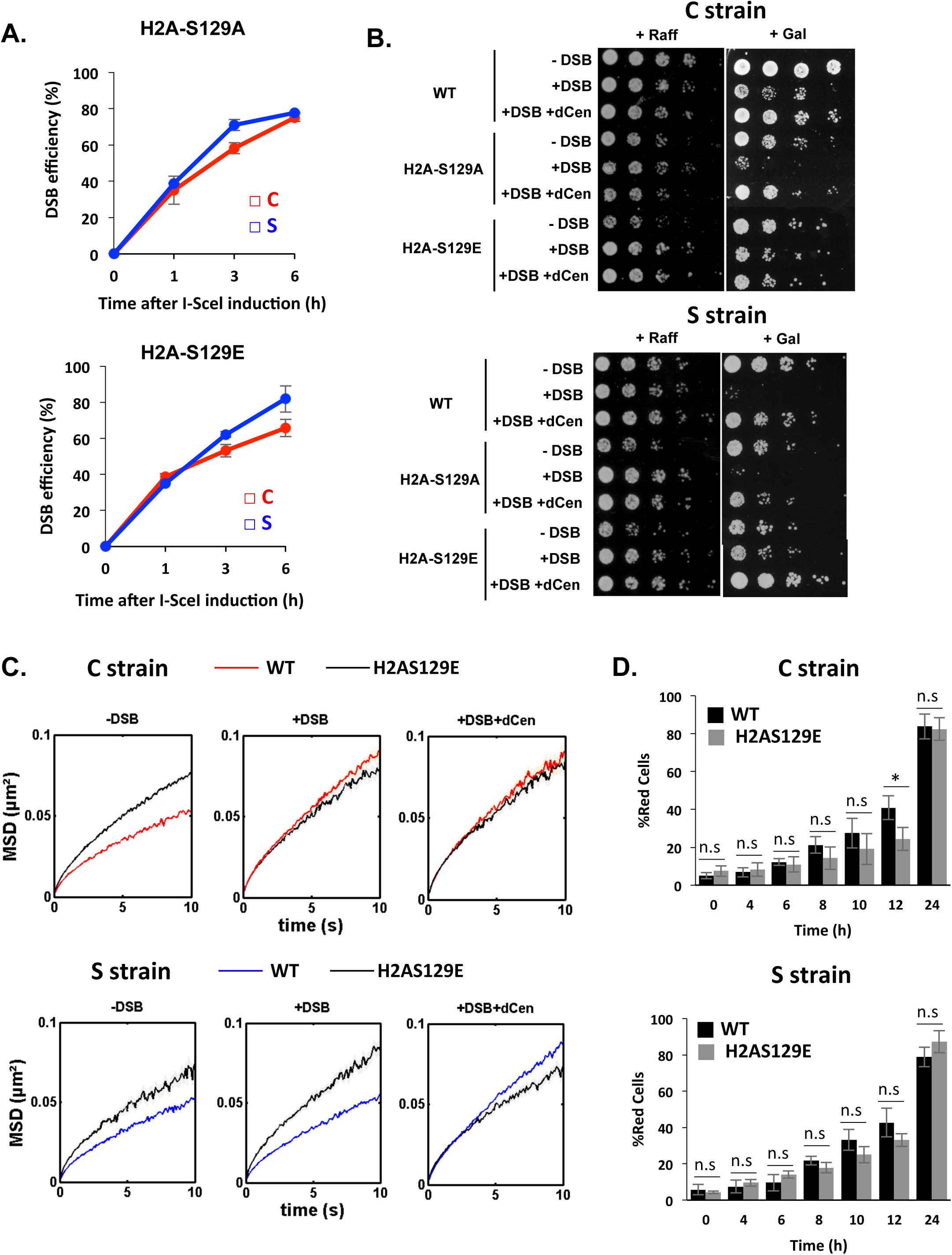
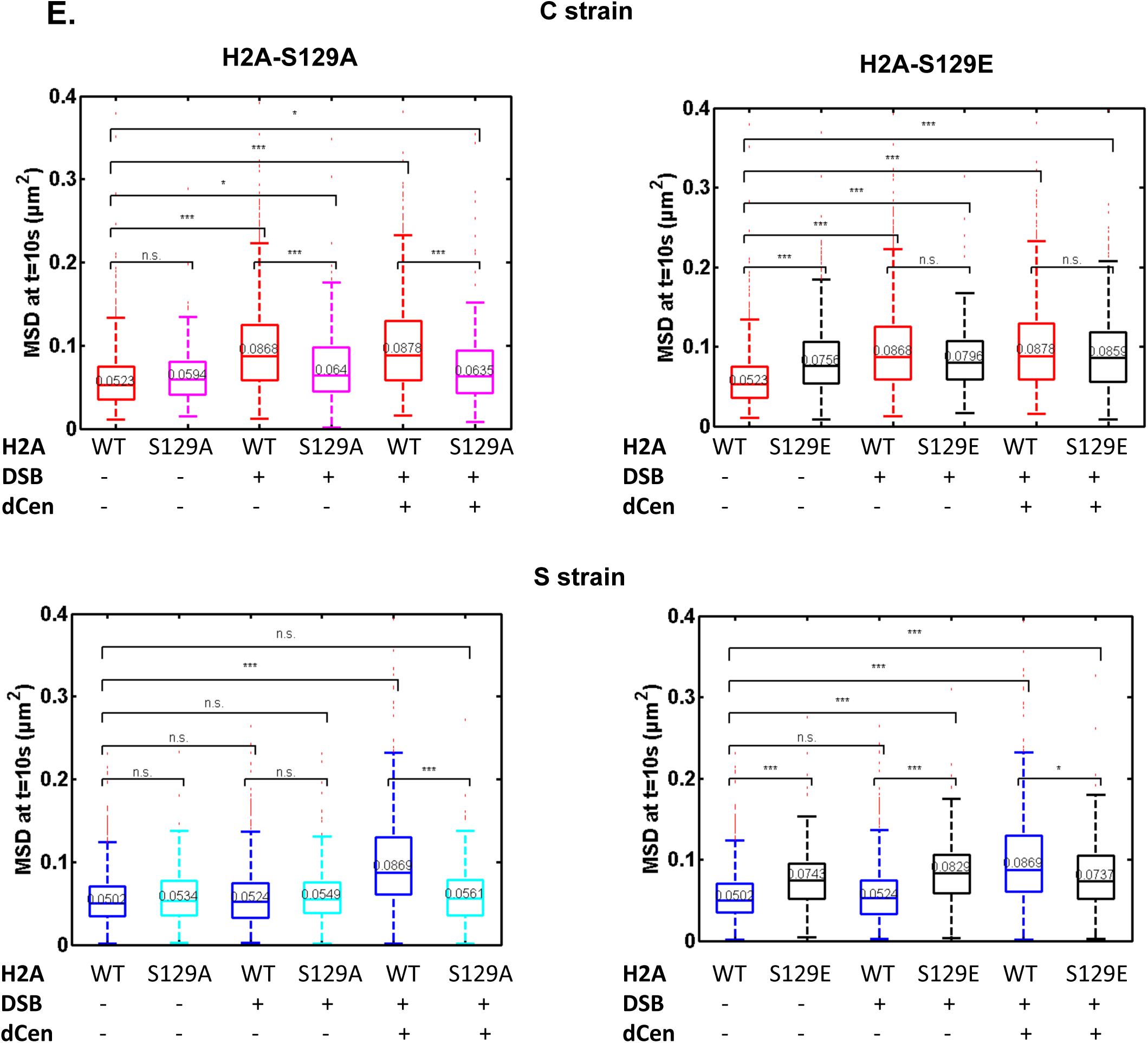
**A**. I-*Sce*I cleavage efficiency is measured in H2A-S129A, H2A-S129E strains by qPCR using primers flanking the I-*Sce*I cutting site. The error bars represent the standard deviation of two independent experiments. **B.** Drop assay (tenfold serial dilutions) showing the sensitivity of WT, H2A-S129A and H2A-S129E strains to long exposition to galactose, inducing DSB in strains C and S, with dCen or empty plasmid. As controls, strains C and S not expressing I-*Sce*I are shown. **C.** MSDs in strains C and S in H2A-S129E mutated backgrounds. MSDs are calculated as in Fig. 2 upon 6h of DSB induction with dCen or empty plasmid (+DSB+dCen, +DSB respectively). Controls without DSB are also shown (- DSB). **D**. HR kinetics upon induction of I-*Sce*I in the presence of dCen in strains C or S on WT (black) and H2A-S129E (grey) background were measured by FACS as in Fig. 1F. Error bars represent the standard deviation of three independent experiments. **E.** Boxplots of the distribution of MSDs at 10s from Fig. 4C and Supp. Fig. 4C, for strains C and S in WT and H2A-S129 mutants without or with a DSB (-DSB, +DSB), with dCen or empty plasmid (+dCen, -dCen). Median values, lower and upper quartiles are shown. Whiskers indicate the full range of measured values, except for outliers represented by small red dots. Parentheses indicate the result of a Wilcoxon rank-sum test between distributions, with the p-value. n.s. non significant, * (p<0.05), ** (p<0.001), *** (p<0.0001).

## Notes

### Competing Interest Statement

The authors have declared no competing interest.

